# Ohmic analogies and metaphorical circuits: Vascular partitioning of leaf air spaces and stomatal patchiness can create apparent undersaturation and gradient inversion

**DOI:** 10.1101/2024.05.30.596638

**Authors:** Fulton E. Rockwell

**Affiliations:** Department of Organismic and Evolutionary Biology, Harvard University, Cambridge MA 02138, USA

**Keywords:** amphistomatous, mesophyll conductance, intercellular air space, stomatal conductance, leaf gas, exchange, Ohm’s, law analogies, patchy stomata, undersaturation, *CO*_*2*_, assimilation

## Abstract

**Rationale:** Analyses of leaf gas exchange rely on an Ohmic analogy that arrays single stomatal, internal air space, and mesophyll conductances in series. Such models underlie inferences of mesophyll conductance and the relative humidity of leaf airspaces, reported to fall as low as 80%. An unresolved question is whether such Ohmic models are biased with respect to real leaves, whose internal air spaces are chambered at various scales by vasculature.

**Description:** To test whether undersaturation could emerge from modeling artifacts, we compared Ohmic model estimates with true parameter values for a chambered leaf with varying distributions and magnitudes of leaf surface conductance (“patchiness”).

**Key Results:** Distributions of surface conductance can create large biases in gas exchange calculations. Both apparent unsaturation and internal *CO*_*2*_ gradient inversion can be produced by the evolution of particular distributions of stomatal apertures consistent with a decrease in surface conductance, as might occur under increasing vapor pressure deficit.

**Main conclusion:** In gas exchange experiments, the behaviors of derived quantities defined by simple Ohmic models are highly sensitive to the true partitioning of flux and stomatal apertures across leaf surfaces. We need new methods to disentangle model artifacts from real biological responses.

## Introduction

Analyses of gas exchange typically rely on an Ohmic analogy that equates a leaf to a single series of resistors (or conductors) that each represent properties aggregated over a whole leaf. For a leaf in a gas exchange cuvette, the surface conductance defined by measurement of transpiration, the water concentration in the well-mixed cuvette air, and the saturated water concentration at leaf temperature, is thought to represent the aggregated conductance of all stomata on the leaf surface in series with a boundary layer (with minor contributions from cuticle). For *CO*_*2*_, this surface conductance is placed in series with a conductance of the intercellular airspace (*IAS*) that comprises the path from the sites of evaporation to the cell walls of the photosynthetic mesophyll. A mesophyll conductance that represents the liquid phase’s path from the cell wall to the chloroplast completes the last of the series (Rockwell et al., 2022). Implicit in this model is the idea that a water or *CO*_*2*_ molecule has access to the entire intercellular airspace regardless of which stomatal pore it travels through.

Yet it is easy to see that this cannot always be the case. Leaves are at some scale subdivided by vasculature into compartments with little lateral gas flow between them (Fig. **1a**,**b**). If stomatal apertures are inhomogeneous across a leaf surface at the areole scale it is possible that most of the flux of gases could occur across some fraction of the leaf surface. In this case, the flux only sees a fraction of intercellular air space conductance, and not the totality of it. In thinking about this effect, it is helpful to recall that resistances arrayed in series are simply summed to find a total resistance, whereas for resistances arrayed in parallel the total effective resistance is the reciprocal of the sum of the reciprocals of the individual resistances. It follows that conductances (the reciprocals of resistances) arrayed in parallel may be simply summed to find a total conductance, but the total effective conductance for conductors arrayed in series is the reciprocal of the sum of the reciprocals of the individual conductances. While the proper ordering of summation operations may appear to be merely a technical issue, in this analysis we will show that a simple Ohmic analogy that treats the internal air space of a leaf as single compartment can, when combined with particular patterns of heterogeneous stomatal apertures, produce behavior that is currently thought to be explicable only if leaf airspaces reach highly unsaturated relative humidities, as low as 80-85% (Cernusak et al., 2018; Wong et al., 2022). These difficulties are closely related to those raised by the effects of “patchy” stomatal behavior on inferences of mesophyll conductances (Laisk, 1983; Downton *et al*., 1988; Rockwell *et al*., 2022).

The classic test for undersaturation involves feeding an inert gas across a leaf to measure the total diffusive resistance, conceptually defined as the resistances of the upper and lower surface (including stomatal, boundary layer and cuticular resistances) arranged in series with the resistance of the intercellular airspace through the thickness of the mesophyll (Jarvis & Slatyer 1970; Farquhar & Raschke 1978, Wong *et al*., 2022). The total resistance that *would* be experienced by water *if* it diffused from one side of a leaf to the other can be calculated (*R*_*tot*_) by correcting for the differences in diffusivities between water vapor and the measured gas. Subtracting from *R*_*tot*_ the two surface resistances seen by transpiration (*R*_*u*_ and *R*_*l*_ respectively) yields an estimate of the difference between any resistance seen by the inert gas but not water, and any resistance seen by water but not the inert gas. Assuming the majority of evaporation occurs on the surfaces of the sub-stomatal cavity (SSC) (Rockwell *et al*., 2022), the former resistance that is particular to the inert gas corresponds to the intercellular airspace (*R*_*ias*_) between the upper and lower SSCs, and the latter resistance particular to water is described alternatively as a cell wall resistance between the symplast and gas phase (Jarvis & Slatyer, 1970; Farquhar & Raschke 1978) or undersaturation of the substomatal cavity (*R*_*unsat*_; Wong *et al*., 2022). An argument can then be made that the *R*_*ias*_ is a physical quantity that is unlikely to change appreciably as the leaf experiences higher vapor pressure differences from ambient air (VPD), and therefore that changes in the difference *R*_*ias*_ - *R*_*unsat*_ represents the development or decay of *R*_*unsat*_ alone.

Complicating interpretations of changes in the difference *R*_*ias*_ - *R*_*unsat*_ is that the subdivision of leaves into areoles has even greater consequences for the transit of inert gases across amphistomatous leaves than it does for water vapor or *CO*_*2*_. To find the total resistance presented by the leaf for the case of an inert gas diffusing from one side of a leaf to the other, the surface and airspace resistances of each individual areole should be summed first to find the total resistance of each areole. These total areole resistances are then aggregated to a whole leaf resistance by taking the reciprocal of the sum of their reciprocals (Fig. **1c**). Although gas exchange systems correctly aggregate the total resistances of the upper and lower surfaces, a simple Ohmic model of a leaf as a single compartment (Fig. **1d**) places these aggregated resistances in series with an effective aggregated resistance for the entire intercellular airspace that fails to respect the subdivisions created by the vasculature: simply subtracting the surface resistances from this model’s definition of total resistance (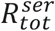, Fig. 1d) will not yield a consistent estimate of *R*_*ias*_ as the distribution of stomatal apertures varies across the upper and lower surfaces. Physically, the problem is that the largest resistance in series dominates the total resistance, masking any low resistance. As a result, asymmetry in stomatal apertures between the upper and lower surfaces of an areole means that the aggregated surface resistances seen by water (*R*_*u*_ and *R*_*l*_) underestimate the actual areole-by-areole contributions of surface resistance to the *R*_*tot*_ actually experienced by an inert gas (Fig. **1C**), leading to an overestimate of the difference *R*_*ias*_ - *R*_*unsat*_.

**Figure 1.**
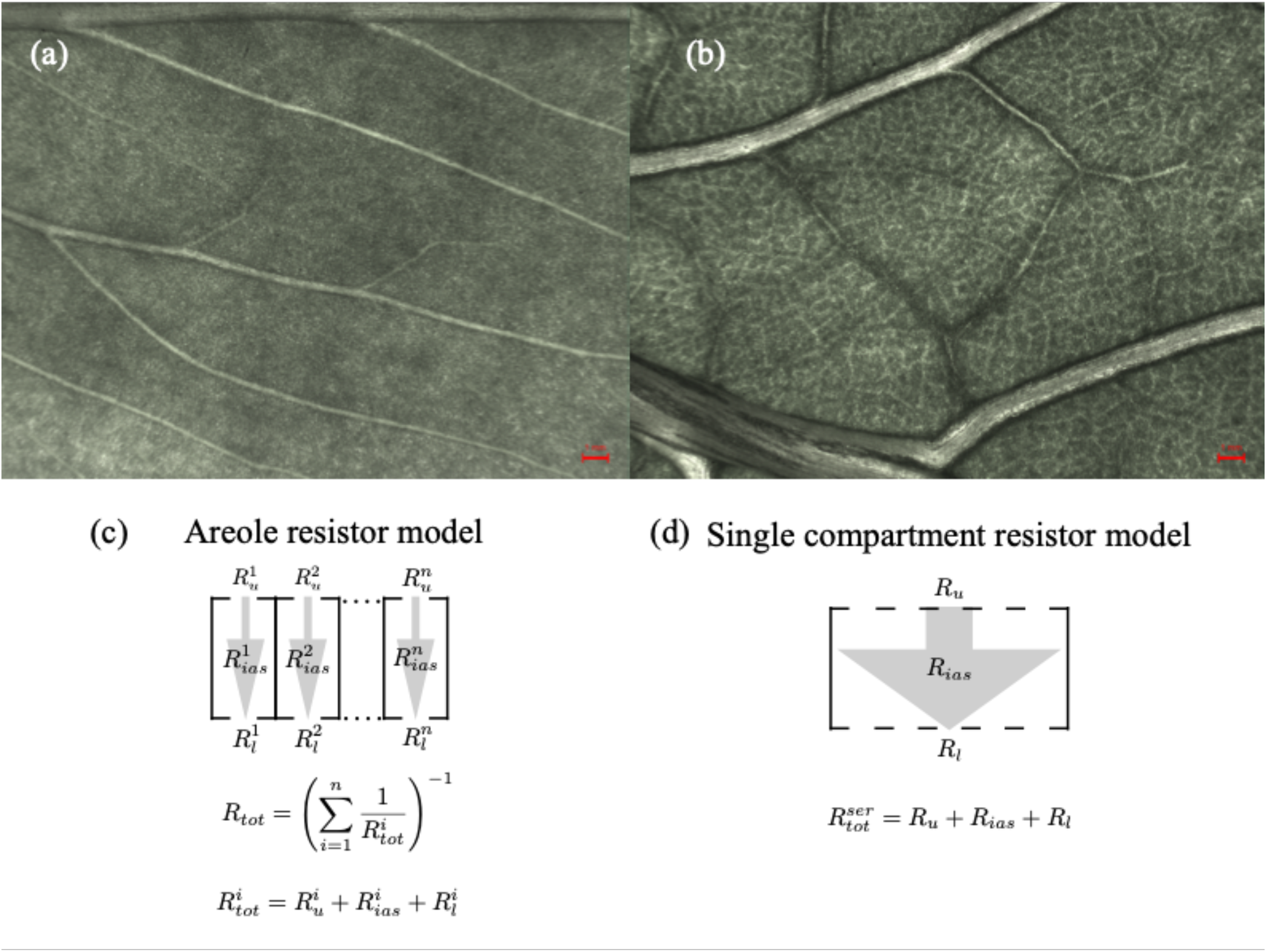
Ohmic models for gas exchange analysis of leaves and the partitioning of leaf air space by vasculature. (**a**) *Vicia faba*, a homobaric leaf with vascular divisions of the mesophyll at the >1 mm scale forming at least six isolated compartments in a total area leaf of 5 cm^2^. (**b**) *Solanum lycopersicum*, heterobaric, with compartmentation at the <1 mm scale and hundreds of areoles within the 5 cm^2^ in view. (**c**) The “true” resistor model of a leaf as areoles arrayed in parallel, each with their own surface and intercellular airspace (IAS) resistances arrayed in series. (**d**) The simplified single compartment model of a leaf typical of gas exchange measurement systems and analyses, as a single array of surface and intercellular airspace resistances in series.

Wong et. al (2022) introduced a new twist to the canonical test for undersaturation comparing inert gas and water resistances discussed so far. In their experiment, the *CO*_*2*_ in the lower cuvette was incrementally lowered until net assimilation from the lower cuvette dropped to zero. This condition then allowed them to solve for the internal differences in *CO*_*2*_ between the inner epidermal surfaces, Δ*c*_*i*_, and calculate an effective resistance of the intercellular airspace to *CO*_*2*_ *R*_*ias,c*_ (Parkhurst *et al*., 1988). By again arguing that this resistance should be invariant to changes in VPD, Wong *et al*. (2022) were able to show that Δ*c*_*i*_ at high levels of VPD deviated from the expected values based on the *R*_*ias,c*_ calculated from low VPD measurements. By then correcting Δ*c*_*i*_ to the expected value, these authors generated an estimate of the degree of unsaturation in the SSC, and so fixed the value of *R*_*unsat*_. As this method does not involve the inert gas, it appears to offer independent confirmation of undersaturation in response to high VPD.

In a previous study, we investigated how the proportion of leaf area active in gas exchange (i.e., “patchiness” in open vs closed stomata, or dead vs alive leaf regions) could, in certain special cases, create the appearance of undersaturation (Rockwell *et al*., 2022). Here we focus on the impact of less extreme forms of “patchiness” in stomatal apertures, adopting the term “compartment” to describe leaf regions composed of areoles with similar surface resistances, and, as noted above, observe the constraint that lateral gas diffusion between compartments is blocked by vasculature. We expect that such a ‘two compartment’ model maps in an unbiased way to a leaf with stomatal apertures clustered around two distinct values, in the same way that a single compartment ohmic model of a leaf maps in an unbiased way to a nearly uniform distribution of stomata apertures (Laisk, 1983). We then consider amphistomatous leaves with two transpiring surfaces, and first investigate how a single compartment resistor model interacts with inhomogeneity across and asymmetry between the surface resistances of a two-compartment leaf to introduce artifacts into estimates of *R*_*ias*_. We then solve for the distributions of *CO*_*2*_ in an amphistomatous two-compartment leaf with distinct fluxes at both surfaces, and simulate experiments of the type in Wong *et al*. (2022) to explore how a single compartment gas exchange model can introduce model dependent artifacts into *CO*_*2*_-based inferences of unsaturation.

## Model Descriptions

### The relationship between resistances in a two-compartment leaf and inferences of R_ias_ using a single compartment resistor model

Standard gas exchange calculations represent a leaf as a linear circuit of aggregated resistances in series (Jarvis & Slatyer, 1970). For an inert gas diffusing from one side of a leaf to the other, this conception results in,

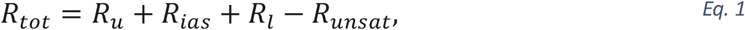

Where *R*_*unsat*_ is a resistance located between the symplast and the substomatal cavity that is seen by transpired water, but not a gas transiting a leaf between the stomata and IAS (Wong *et al*., 2022; note that, for the purposes of comparison, all resistances are defined as the equivalent resistances that would be experienced by water vapor). At low VPD, *R*_*unsat*_ is assumed to be zero, and *eq*.*1* can then be re-arranged to define the unknown resistance of the intercellular airspace in terms of the measured surface and total resistances for a leaf; we write this inference of *R*_*ias*_ as,

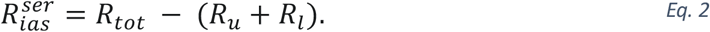

We next consider a leaf with two-compartments that may vary in size between measurement conditions; exploratory simulations with larger numbers of compartments (up to ten) did not appear to introduce any qualitatively new behavior that might have justified the additional complexity. With α_i_ the leaf area fraction of and *R*^*i*^ a resistance associated with the *i*^*th*^ areole (Fig. **1**), this description yields for two compartments (labelled 1, 2),

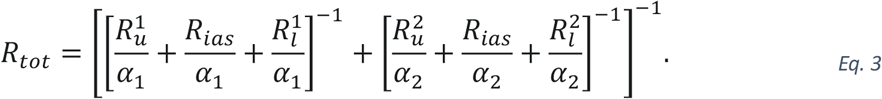

The total conductances of the upper and lower surfaces are similarly defined as,

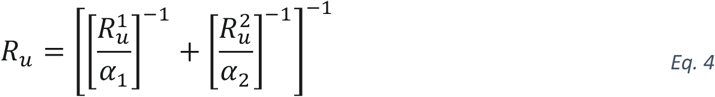

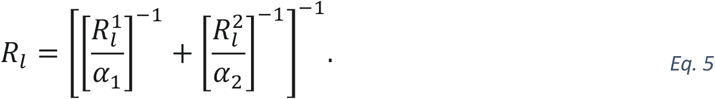

In the case that the *CO*_*2*_ concentration of the lower cuvette has been reduced to the point that there is no net flux of *CO*_*2*_ across the lower leaf surface, the total resistance seen by *CO*_*2*_ (again, re-scaled to the resistance seen by water vapor) has the form,

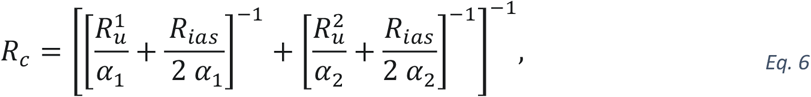

where the factor of two enters as, in the limit of homogenous assimilation through the leaf thickness, the average *CO*_*2*_ molecule only travels half the leaf thickness (Parkhurst *et al*., 1988). An effective resistance for the *IAS* may then be defined by a single compartment model as,

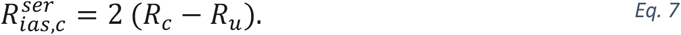

### Gradients of CO_2_ in a two-compartment gas exchange model of an amphistomatous leaf

Parkhurst *et al*. (1988) presented solutions for the distribution of *c*_*i*_ in a single compartment hypostomatous leaf, and for an amphistomatous leaf in the special case that the fluxes from the upper and lower surface were symmetrical. Although Wong *et al*. (2022) performed gas exchange experiments on amphistomatous leaves, the experimental condition that the net *CO*_*2*_ flux from the lower cuvette to the leaf was zero justified using Parkhurst’s hypostomatous model to solve for *c*_*i*_. Here, to test the inferences of a single compartment Ohmic model of leaf gas exchange, we solve simultaneously for the internal gradients and fluxes associated with each compartment in a two compartment leaf model that allows for inhomogeneous stomatal apertures across the leaf surfaces, and the general case of asymmetric fluxes of *CO*_*2*_ across the upper and lower surfaces. In the relations that follow, for the sake of clarity we neglect details of cuticular conductance, ternary corrections, and corrections for the differences between the effective diffusivities for water vapor and *CO*_*2*_ in the leaf boundary layer. We then express the total surface conductances (stomatal and boundary layer) with respect to water vapor, correcting for *CO*_*2*_ where appropriate by dividing by the ratio of water to *CO*_*2*_ diffusivities (1.6; Wong *et al*., 2022). The internal mole fraction (mol *CO*_*2*_/mol air) gradient of *CO*_*2*_ for a single areole of an amphistomatous leaf is then found to be (see Supporting Information Notes S1 for a derivation),

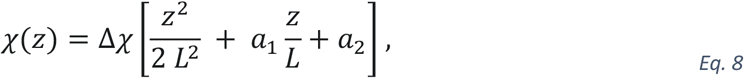

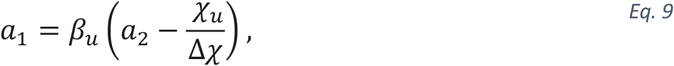

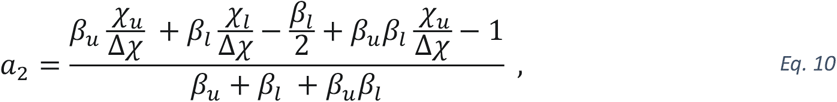

and,

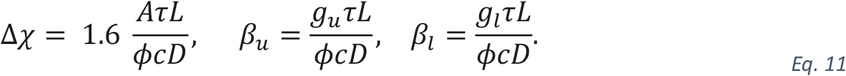

With *χ*_u_, the mole fraction for the upper cuvette, and the compartment specific quantities g_u_, and g_l_, the conductances of the upper and lower surfaces to water vapor, *A* the assimilation rate, and τ the tortuosity, the only remaining unknown is *χ*_*l*_, the *CO*_*2*_ mol fraction for the lower cuvette that renders the net assimilation rate from the lower cuvette zero. For two compartments with area fractions α, this value is then fixed by evaluating the above equations for each compartment and setting their combined fluxes at the lower surface equal to zero:

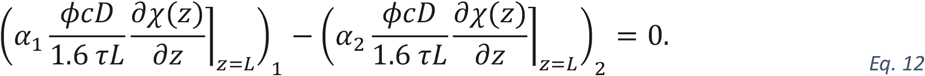

This last equation reminds us that only the *net* flux between the lower cuvette and a leaf is zeroed in the procedure of Wong *et al*. (2022), and in general we should expect that offsetting fluxes exist at the areole scale. Once *χ*_*l*_ is known, *eqs*. 8-11 can be evaluated to find the values of *CO*_*2*_ mol fraction inside the lower and upper epidermal surfaces as *χ* (*L*) and *χ* (*0*) respectively for each areole. These values provide estimates of the Δ*χ* within each areole that may then be compared to that of a single compartment model of a leaf. Assuming saturation of the airspaces (i.e., at low VPD),

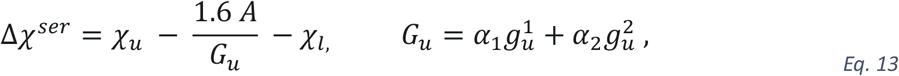

where *G*_*u*_ is the total conductance of the upper surface to water vapor as reported by a gas exchange system. Wong *et al*. (2022) then define a corrected Δ*χ*^*ser*^ for high VPD conditions, under which undersaturation could lead to errors in *G*_*u*_, by arguing that the conductance of the internal airspaces to gaseous diffusion should be relatively constant and the ratio Δ*χ*^*ser*^/*A* should therefore be independent of VPD:

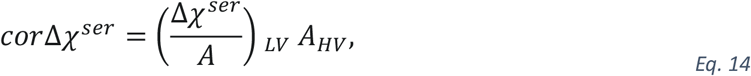

where *LV* and *HV* refer to values seen at low and high VPDs respectively. In the single compartment model, the mol fraction inside the lower leaf surface is assumed to be in equilibrium with the lower cuvette, Δ*χ* (*L*) = Δ*χ*_*l*_, and a corrected estimate for the top of the mesophyll 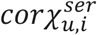 follows as the value in the lower cuvette plus the corrected delta within the leaf,

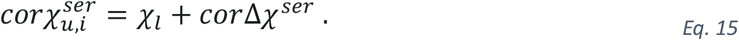

Next, a corrected upper surface conductance and mole fraction of water vapor in the upper substomatal cavity follows as,

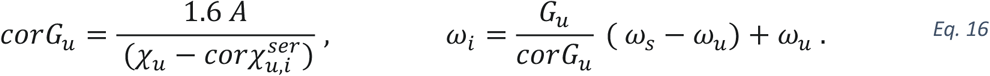

Assuming the same degree of undersaturation exists in the lower SSC, the resistance associated with undersaturation (*ω*_*i*_ ≪ *ω*_*s*_) can be found from the differences between the corrected and uncorrected surface conductances,

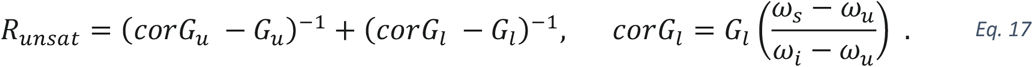

This definition includes the definitions of the total surface resistance and its corrected form,

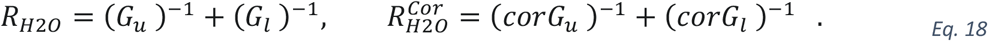

An estimate of *R*_*ias*_ corrected for undersaturation follows as,

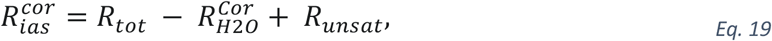

Where it may be helpful to recall that *R*_*tot*_ is an experimentally measured value based on the diffusion of an inert gas through the whole leaf.

### Simulations of a sequence of gas exchange experiments to produce apparent undersaturation

To test whether a two-compartment model that remains near saturation could produce apparent undersaturation and *c*_*i*_ gradient inversion, we constructed a series of leaf states consisting of varying surface resistances and area fractions for each compartment, and a single conserved intercellular airspace resistance for the leaf. We then iteratively varied the assimilation rate for each compartment to produce a consistent but fictitious relationship between *A* and the *c*_*i*_ at the upper surface (Supplemental Fig. S1), as in a typical gas exchange *A*/*c*_*i*_ curve, similar to the values given in Table 1 of Wong *et al*. (2022). We did not attempt to model the experiment depicted in Figure 3 of that study as the values were not included in the supplemental data.

Modeling experiments based on the reported data, while possible, proved cumbersome due to the difficulty of iteratively matching experimental observations to ‘true’ leaf states by varying eight unknown parameters (e.g., assimilation rate, the conductance for each surface and compartment, area fraction, the true *A*/ci relationship).

## Results

### Analysis of errors arising from single compartment models of a two-compartment leaf

We first focus on how well the single compartment model (Fig. **1d**) estimates *R*_*ias*_ based on measurements of the aggregated total surface resistances (seen by water) and total transport resistance across a leaf (as seen by an inert gas, eqs. 2,3,4,5, or *CO*_*2*_, eqs. 4,6,7) in a hypothetical two-compartment leaf. For the null case that stomatal apertures are homogeneous between compartments on each leaf surface, and symmetric *within* each compartment between leaf surfaces (i.e., 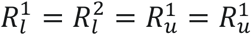 in a two-compartment model), the single compartment model for transport both across the whole leaf (inert gas), as well as from the upper surface to half the distance through the IAS (*CO*_*2*_), proves robust (Fig. **2a**). For the case of asymmetric but homogeneous surface resistances (i.e., 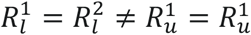), the single compartment model again performs well, as homogeneity renders differences in the order of summations irrelevant. Errors start to emerge when surface resistances are inhomogeneous between compartments on a a surface, though still symmetric between surfaces. This effect occurs as high values of symmetric surface resistances effectively mask the low resistance of the IAS between them. In the single compartment series model, the resistance of the IAS is that of the entire IAS (eq. 1), which underestimates the resistance of the relevant subsection of the IAS that is between the low resistance surfaces: that is, the single compartment model uses *R*_*ias*_ when it would be better (to a first approximation) to use *R*_*ias*_*/α* (i.e., the dominant term in eq. 3, where α is the area fraction of the low resistance compartment). Notably this error effects both the single compartment model for the total resistance to transport across the leaf (eq. 1) and the transport of *CO*_*2*_ through the upper surface and halfway through the IAS (eq. 6) identically, leading to identical overestimates of *R*_*ias*_ (eq. 2; eq. 7).

Adding asymmetry to inhomogeneity breaks the symmetry in errors for *CO*_*2*_ and inert gas transport. The estimate of *R*_*ias*_ based on *CO*_*2*_ is identical to the symmetric homogenous case as here we have assumed that there is no transport across the lower surface, and so asymmetry does enter the calculations. A very different outcome occurs for the estimate based on inert gas transport across the leaf, as here the effects of asymmetry are strong, leading to a five-fold increase in the error (Fig. **2a**; 8.4 vs 1.6). Asymmetry results in the high resistance surfaces in a two-compartment leaf masking the low resistance surfaces (eq. 3), leading to large errors by the single compartment model (eqs. 1 and 2).

The pattern of single compartment model failure is broadly similar when one considers the same distributions of resistance in a gas exchange model (Fig. **2b**). In this more complete description, for a two compartment leaf, the expectation of equilibrium across the lower leaf surface when there is zero net assimilation of *CO*_*2*_ from the lower cuvette fails: the constraint of zero net assimilation means that the fluxes between the two individual compartments and the lower cuvette will be equal in magnitude and opposite in direction, and only each individually zero for the case that the distribution of resistances in each compartment is identical. The existence of such off-setting fluxes results in the single compartment calculation over-estimating the change in *CO*_*2*_ concentration across the mesophyll (eq. 13), and so overestimating *R*_*ias*_ (Fig. **2b**).

In the inhomogeneous and asymmetric case 4, the symmetry between the behavior of the resistor (Fig. **2a**) and gas exchange models (Fig. **2b**) breaks down. In the resistor models, the *CO*_*2*_ based inference has a certain artificiality – it shows the effect of assuming a single compartment, but it is not a calculation that could ever be made from data, as one can see by inspection of eqs. 6 and 7 it requires knowledge of the true value of *R*_*ias*._ In the full gas exchange model, the *CO*_*2*_ based estimate of *R*_*ias*_ is determined by experiment and does not require prior knowledge of the true value. This comes at the ‘price’ of having to assume that the *CO*_*2*_ flux across the lower surface is zero, making the estimate sensitive to the distribution of resistances on the lower surface (Fig. **2a**,**b**). The error therefore converges with that of the inert gas inference (Fig. **2b**). All of the calculations in Fig. 2 assume that saturation of the leaf air space holds, and so at least under experimental conditions where saturation is expected (i.e., low VPD), these simulated gas exchange data suggest that, in real experiments, we should expect to see similar errors in *R*_*ias*_ for both *CO*_*2*_ and inert gases.

**Figure 2.**
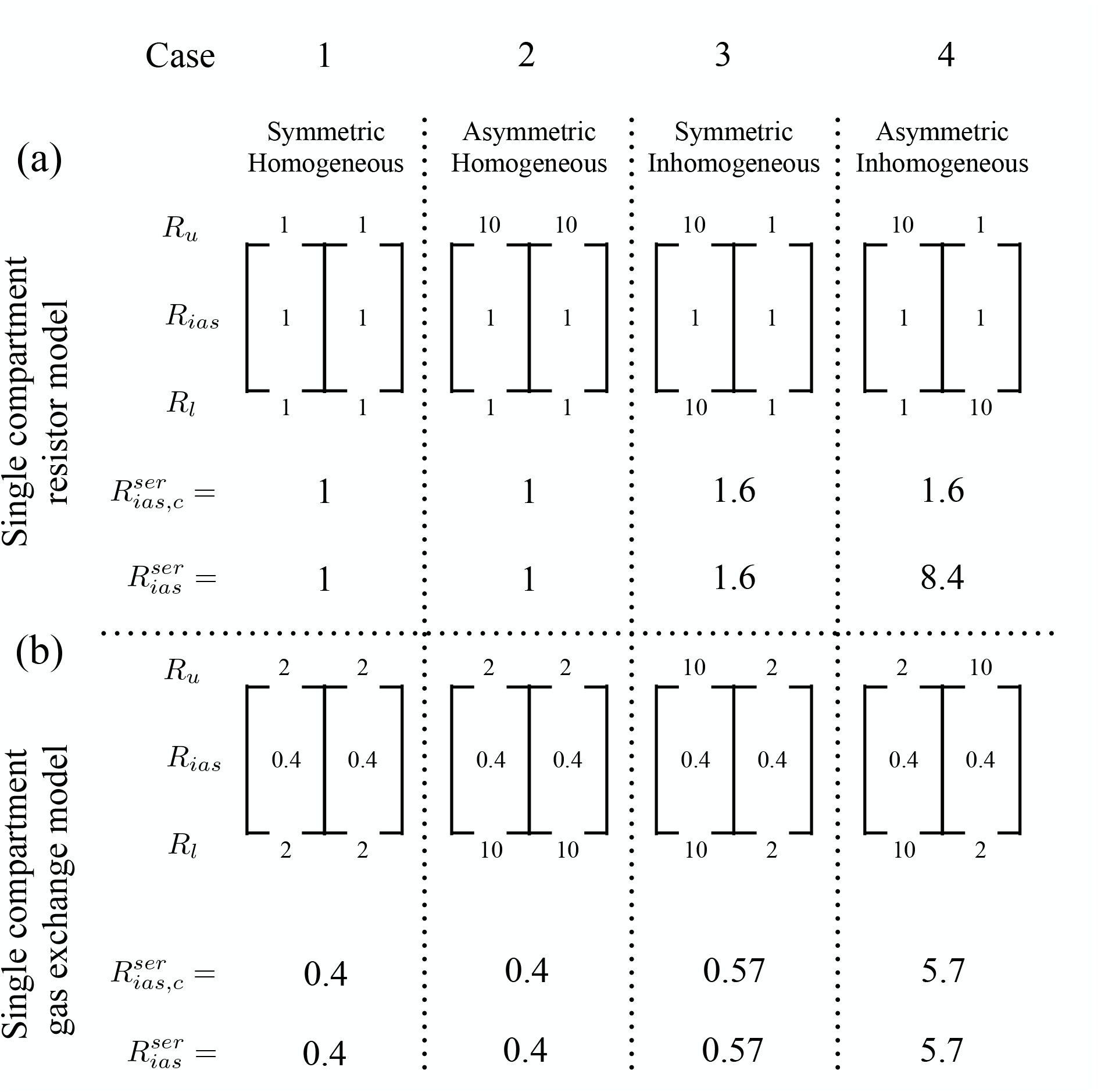
Diagrams presenting the actual resistances of the upper (*Ru*) and lower (*R*_*l*_) surfaces and intercellular air space (*R*_*ias*_) for a leaf with its internal airspace divided into two compartments. The corresponding estimates of *R*_*ias*_ inferred from *CO*_*2*_ or an inert gas are shown below each case. (**a**) Inhomogeneity in resistances across both lower and upper surfaces lead to errors in single compartment model estimates of IAS resistance from the perspective of both *CO*_*2*_ fluxes (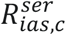 ; eq. 6) and inert gas transport (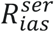 ; eq. 2) across the leaf (cases 3,4). *CO*_*2*_ transport in the resistor model is insensitive to asymmetry by construction (due to zero lower flux), even as for the inert gas asymmetry has a large impact on the effective resistance of each compartment (cases 3 versus 4). (**b**) For a single compartment model of the actual gas exchange fluxes, *CO*_*2*_ based inferences of *R*_*ias*_ track those of an inert gas as the individual fluxes of *CO*_*2*_ from the lower cuvette to each compartment are equal and opposite but non-zero, making 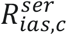 sensitive to asymmetry between leaf surfaces.

**Figure 3.**
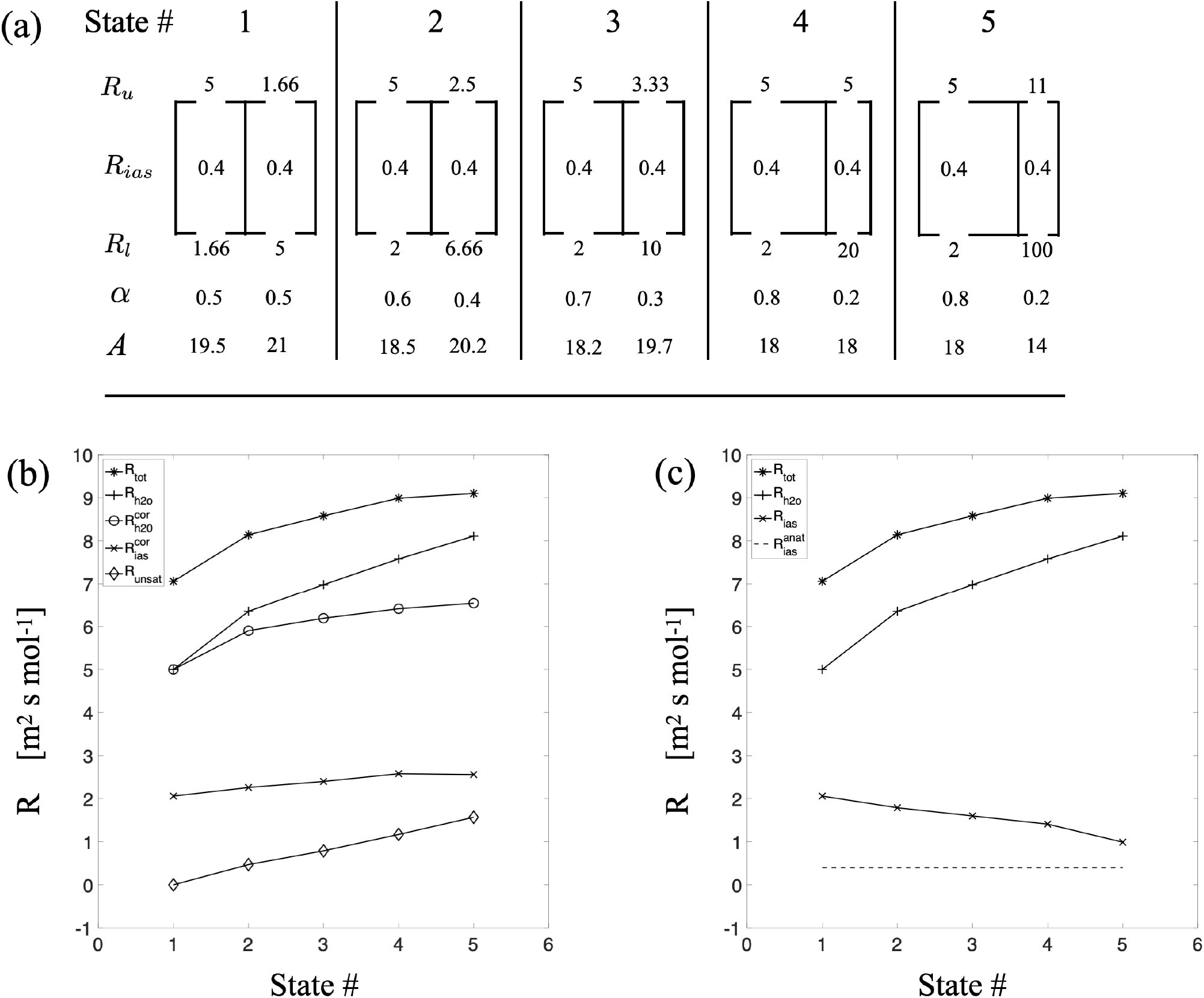
Model bias over a series of surface resistance states (e.g., stomatal VPD responses) can reproduce the growth of an unsaturation resistance observed by Wong et al. (2022). (**a**) Diagram of the true surface and intercellular airspace resistances, area fractions (α) associated with each mode (i.e., compartment) of a bimodal distribution of resistances, and assimilation rates (*A*) for each level of VPD. Surface resistances begin at low VPD in an asymmetric inhomogeneous state characterized by a large overestimate of *R*_*ias*_. As VPD increases, the surface resistances shift such that the amount of leaf area (number of areoles) associated with high surface resistances shrinks (lower α) even as the magnitude of the resistances increases. (**b**) Evolution of resistances over the above series of states, showing the convergence of *R*_*h2o*_ with *R*_*tot*_ and *R*_*unsat*_ with *R*_*ias*_ that has been interpreted as evidence of unsaturation. In fact, forcing the conservation of Δ*c*_*i*_/A across the series of states leads to near conservation of 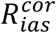 at an artifactually high level, and the growing appearance of an unsaturation resistance. (**c**) Uncorrected data: the decline in apparent *R*_*ias*_ and the convergence of *R*_*h2o*_ toward *R*_*tot*_ over the sequence of states, thought to be non-physical, in fact represents convergence of estimated *R*_*ias*_ with the true anatomical value. Convergence occurs due to a reduction in the effects of asymmetry and inhomogeneity on inferred *R*_*ias*_ as inhomogeneity across the upper surface at first declines from state 1 to 4, and then is restricted to a small amount of leaf area in state 5.

In early work on undersaturation in leaf airspaces (Jarvis & Slatyer, 1970), the total resistance seen by an inert gas and the surface resistances defined by transpiration fixed a value for *R*_*ias*_ by assuming that at low VPD *R*_*unsat*_ was zero (eqs. 1,2;), and then as VPD was increased the value of *R*_*ias*_ - *R*_*unsat*_ was observed to decline: assuming *R*_*ias*_ remained unchanged, this implied the growth of *R*_*unsat*_ between the SSC (the presumed cite of *w*_*i*_) and a location where an assumption of saturation held. Given that the errors discussed so far lead to overestimates of *R*_*ias*_, for the single compartment model to create the appearance of undersaturation requires that the errors be largest at low VPD (asymmetric, inhomogeneous), and then decline due to greater symmetry, or greater homogeneity, or both as VPD increases. If this is the case, then one would predict that other drivers of stomatal movements aside from VPD should create apparent changes in *R*_*ias*_ - *R*_*unsat*_. Some evidence for this exists: Farquhar and Raschke (1978: Fig. 1,2) show time courses of stomatal opening in light accompanied by variations in *R*_*ias*_ - *R*_*unsat*_.

### The evolution of apparent undersaturation through a sequence of surface resistance states

The experiments of Wong *et al*. (2022) introduced a second method for detecting *R*_*unsat*_, independent of the transport resistance seen by an inert gas. If *R*_*ias*_ is again assumed not to change, then the ratio of Δ*c*_*i*_ across the mesophyll to assimilation *A* should be fixed. Deviations from the expected Δ*c*_*i*_ provide a basis for correcting the upper surface resistance and estimating *R*_*unsat*_ (eqs. 14-17). Here we show that it is possible to construct a sequence of leaf surface resistance states characterized by increasing stomatal resistance, as might occur in response to increasing levels of VPD, which reproduce these patterns in a leaf for which the saturation assumption holds by hypothesis (Fig. **3a**). The leaf starts in an asymmetric and inhomogeneous state, characterized by a five-fold overestimate of *R*_*ias*_ (State 1; Fig. **3a**,**b**). The subsequent evolution of states is characterized by little change in the resistances of compartment 1 as compartment 2 undergoes stomatal closure, with in an increase in inhomogeneity for the lower surface (even as homogeneity increases on the upper surface until the transition from state 4 to 5) and little change in asymmetry. While this trajectory might naively be expected to bias apparent *R*_*ias*_ to even higher values, this tendency is counteracted in part by an increasing area fraction of compartment 1 from 0.5 to 0.8 of the leaf surface area as *R*_*H2O*_ increases in compartment 2. The net effect is that *R*_*H2O*_ approaches *R*_*tot*_ for the inert gas even as *R*_*ias*_ drops toward its true anatomical value (Fig **3c**).

Ironically, the reduction of error in *R*_*ias*_ leads to the introduction of error when the corrections prescribed by Wong *et al*. (2022) and based on the conservation of internal Δ*c*_*i*_/A (eq. 14-19) are applied. This error arises due to the evolution of a small, high resistance compartment that is nearly isolated from the lower cuvette, such that the assumption of *CO*_*2*_ equilibrium across the lower leaf surface fails. As the correction procedure has the overestimate of *R*_*ias*_ from the initial state baked into it (eq. 14), application of the correction across later states results in a near constant but artifactual ‘corrected *R*_*ias*_’ that parallels the growth of *R*_*unsat*_ (Fig. **3b**). The main point here is that the sequence of leaf states in Fig. **3a** produces a sequence of changes in resistances similar to those reported in Figure 3 of Wong *et al*. (2022), yet without sacrificing the saturation assumption.

### The evolution of an apparent inverted gradient in c_i_

In some experiments at high VPD, Wong *et al*. (2022) note that calculating Δ*c*_*i*_ across the mesophyll based on the saturation assumption for the upper surface, and the assumption of equilibrium with the cuvette for the lower surface, results in a non-physical negative value. Such a result is reproduced here for single compartment model calculations, even as the actual Δ*c*_*i*_ in each compartment remains positive (Fig. **4**). In the initial state, the single compartment estimate of Δ*c*_*i*_ over-estimates the true compartmental values (31.5 vs 4 and 8 PPM respectively; Fig. **4b**,**c**). The error arises as the single compartment model is biased at the upper surface toward the higher *c*_*i*_ of the two compartments, and toward the lower *c*_*i*_ of the two compartments at the lower surface, inflating Δ*c*_*i*_. In the final state 5, the smaller high resistance compartment has uncoupled from the lower cuvette, whose concentration then tracks the larger low-resistance compartment, even as at the upper surface the high resistance of the smaller compartment drives the single compartment model to over-estimate the drop in *CO*_*2*_ across the upper surface. The net result is that the single compartment model calculates a *c*_*i*_ value at the top of the mesophyll below that seen in the lower cuvette (Fig. **4e**,**f**).

Applying the correction procedure based on conservation of Δ*c*_*i*_/*A* (eqs.13-19) results in an increase in single compartment model predicted upper surface *c*_*i*_ from 236 to 266 ppm, dropping RH for the IAS from 100% to 96%, equivalent to a drop in liquid phase water potential from 0 to -5 MPa. This effect in our simulations is then at the low end of effect sizes reported by Wong et al. (2022), although our simulations, perhaps due to the simplicity of a two-compartment model, are sensitive to small changes in the varied parameters: for example, raising the assimilation rate (*A*) of compartment 2 from 14 to 16 μmol m^-2^ s^-1^ doubles the negative Δ*c*_*i*_ from -3 to -6 PPM (data not shown). The important point is that a minimal model respecting the compartmentation of real leaves can be shown to produce, from the perspective of current models of leaf gas exchange, a degree of undersaturation and gradient inversion thought to be impossible for real leaves.

## Discussion

> “A metaphor’s … ambition to illuminate blinds those who
>
> create metaphors. In my distrust of metaphors I feel a
>
> kinship to George Eliot: “We all of us… get our thoughts
>
> entangled in metaphors, and act fatally on the strength of
>
> them.”
>
> Excerpt From: *Dear Friend, from My Life I Write to You in*
>
> *Your Life*, by Yiyun Li.

Analogies are a fundamental tool in science for mapping new phenomena to something already known, and so building new understanding or creating new hypotheses. In plant hydraulics and gas exchange, electrical models provide a useful formalism for analyzing flows and fluxes and defining conductances that may then be related to underlying plant structures as well as chemical and physiological states. As in standard gas exchange calculations the fluxes are normalized to the leaf surface in the cuvette, it is natural to define the proportionalities between fluxes and driving forces similarly, resulting in a single compartment model of a leaf as a simple linear circuit. When we forget that such demonstrably powerful and successful analogies are limited to only a specific set of similarities, and fail to remember that a leaf may not resemble a simple linear circuit in all of its possible states and behaviors, we have come to think the leaf *is* a circuit – the analogy has become a metaphor. The argument of Wong *et al*., (2022) that a negative Δ*c*_*i*_ defined at the whole-leaf level is non-physical commits a metaphorical fallacy in failing to consider that the model itself may not reliably map back to the true diversity of physical properties and behaviors evidenced by a leaf.

Rejecting the argument of non-physicality does not allow us to reject undersaturation as an explanation, it simply means we are not forced to accept it. One might then ask, how plausible is the sequence of states that lead to the rejection of undersaturation? At first glance, the initial state of asymmetric and inhomogeneous stomatal apertures may seem odd. It essentially consists of areoles that are much more open for gas exchange on one surface than the other, and that are then arrayed in parallel (open side up or down) to form a leaf. This would appear to run counter to the hypothesis that the adaptive value of amphistomy is to shorten the path for internal *CO*_*2*_ transport by opening both sides of the leaf (Parkhurst, 1978), an idea that supports an expectation of symmetry. Yet the empirical behavior of stomata and patchiness, well reviewed by Mott and Buckley (2000), support the patterns of surface conductance states adopted here: Stomatal apertures are known to vary widely across a leaf surface (Meyer & Genty, 1998); stomata in the same areole but on opposite sides of the leaf respond to environmental forcing (e.g., light, VPD) independently of the other surface (Mott, 2007); leaves under optimal conditions for photosynthesis still operate far below their theoretical maximum stomatal conductances calculated from anatomy (McElwain et al. 2016); and finally, fluorescence imaging of the amphistomatous *Xanthium strumarium* showed some patches with with closed stomata on both surfaces, some more open on both surfaces, and others one other surface more open than the other (Mott et al., 1993).

What little is known about the dynamics of patches is also broadly supportive. In the simulations employed here, reaching the final state 5 requires that even as the surface resistances in the second compartment increase, the amount of leaf area associated with this high resistance compartment decreases. In defense of this perhaps odd trajectory, Cardon *et al* (1994) showed in *Helianthus annuus* that patches defined as collections of similarly behaving pixels in imaging fluorescence can show a mix of steady and oscillatory behavior with an overall secular trend toward opening or closure at the whole leaf level. From the perspective of two compartments composed of areoles with similar values, such dynamics would appear as changes in the amount of leaf area associated with each compartment over time, as occurs over the sequence of states (1 to 5) in this analysis. How often leaves actually display such bimodal (or highly skewed) distributions of stomatal apertures remains controversial (Meyer & Genty, 1998; Mott, 1995). Here, the important point is that to the extent any distribution of stomatal apertures across a leaf is well-approximated by at least two ‘bins’ of clustered values, a two-compartment model clearly demonstrates the potential for artifacts when the conventional single compartment Ohmic analogy is applied to leaf gas exchange. Even if the two-compartment model fails in some specifics regarding stomatal behavior, it suggests that a *general* recipe for artifacts involves a reduction of asymmetry and inhomogeneity across a leaf coincident with the growth of high resistance over a minority of the leaf area.

In the end, theoretical demonstrations of plausibility cannot substitute for experimental confirmation. The most direct way to check for undersaturation and the partitioning of surface resistance between stomatal and non-stomatal limitation of gas exchange would be to visualize the actual stomatal apertures across a leaf, an approach that suffers from the need to resolve micron scale apertures over centimeter scale surfaces (Kaiser & Kappen, 2000). Fluorescence imaging has been used to investigate stomatal statistics (Meyer & Genty, 1998), but is ill-suited for resolving separate upper and lower surface resistances in a simple way as the fluorescence responses visible for either surface integrate changes in total *CO*_*2*_ delivery (Cardon *et al*., 1994). Leaf replicas offer a way of capturing stomatal level detail across centimeter scale distances on each surface, but the common choice of fast-curing polyvinyl siloxane (i.e., dental impression material) does not provide enough resolution to capture stomatal aperture, and while PDMS can resolve micron scale features curing requires hours, during which time stomata will surely shut (Soffe *et al*., 2019). Fast UV curing elastomers have been used to create replicas of leaf surface for contact angle measurements (Park *et al*., 2012), but their ability to capture stomatal apertures remains untested.

Online carbon and water isotopes provide an alternative method for detecting undersaturation (Cernusak, Holloway-Phillips *et al*., 2019), and have been reported to produce similar results to the gas exchange technique (Wong *et al*., 2022). As isotopic techniques also rely on a single compartment model of leaf gas exchange, it seems likely they are subject to the same types of biases reported here. An entirely different experimental approach, one independent of stomatal states, is provided by a nanogel (aquadust) that can be infiltrated into leaves, where it becomes adsorbed onto the internal leaf surfaces, reporting local water potentials by FRET fluorescence (Jain *et al*., 2019). These experiments support the general phenomena of unsaturation, but, comparing observed minimum relative humidities, perhaps to a lesser degree than reported in the gas exchange and isotopic experiments (Cernusak et al., 2018; Holloway-Phillips *et al*., 2019; Jain et al., 2023; Jain et al., 2024; Wong *et al*., 2022). A second important difference in these studies is that the unsaturation seen by aquadust develops primarily due to water stress on the supply-side; for well-watered plants, demand-side increases in VPD do not elicit unsaturation detectable by this probe. The difficulty of capturing the actual distributions of stomatal apertures across both leaf surfaces makes nanogel-based water potential probes (aquadust), infiltrated into the sites of evaporation, the most practical currently available independent method for verifying undersaturation states.

## Conclusion

Inhomogeneity in stomatal apertures across, and asymmetry between, leaf surfaces can result in mathematical biases in the way that single compartment models calculate transport resistances, and these biases can create both the appearance of undersaturation and *c*_*i*_ gradient inversion. Although errors in estimates of undersaturation have been the focus of the analysis presented here, the biases introduced by single compartment modeling of multi-compartment leaves are of a general nature. All derived quantities calculated from gas exchange fluxes, including mesophyll conductances/resistances and concentrations, are potentially affected. New methods for capturing stomatal heterogeneity, supplemental or integrated into gas exchange equipment, are urgently needed if we are to disentangle model bias from real physiological responses to environmental forcings (e.g., light, *CO*_*2*_, VPD, temperature) when these forcings also affect the distribution of stomatal apertures across a leaf’s surfaces.

## Supporting Information

Additional supporting information may be found in the online version of this article.

**Notes S1**. General solution for the internal CO_2_ gradients across the mesophyll of an amphistomatous leaf.

## Acknowledgements

This work was supported by the National Science Foundation (NSF-MRSEC DMR-2011754), the Air Force (AFOSR FA9550-21-1-0283) and the Star-Friedman Challenge for Promising Scientific Research (Harvard University). N.M. Holbrook and A.D. Stroock and provided many insightful comments on the manuscript and figures, as well as, with their respective lab members, thoughtful discussion of the main ideas and their presentation.

## Competing interests

The author has no competing interests.

## Data availability

Mathematica code (Wolfram Research, Champaign Illinois, USA) used for generating the data in figures 2, 3, and 4 is available from the author.

**Figure 4.**
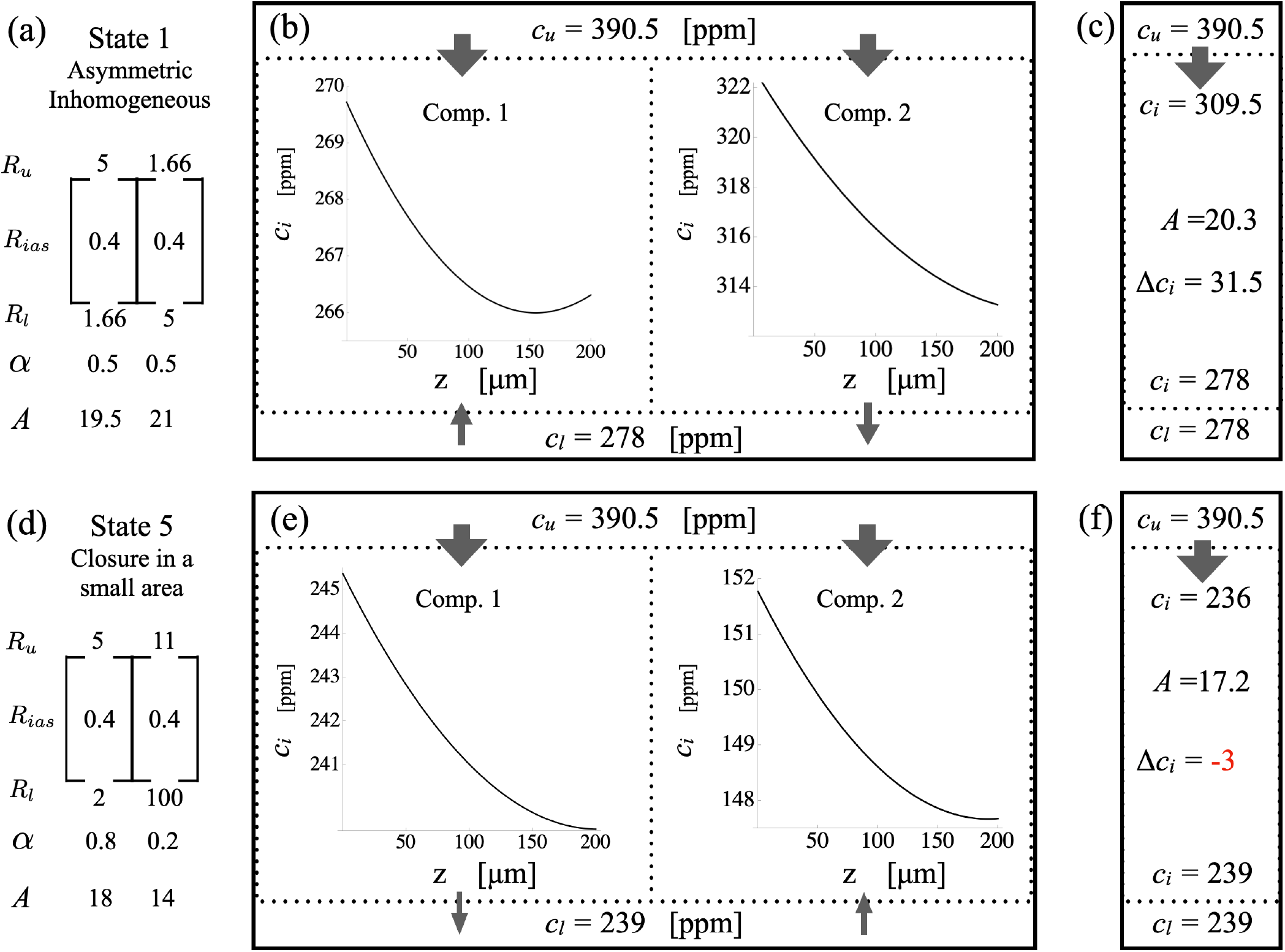
*CO*_*2*_ gradient inversion as an artifact of a single compartment model. For the first and final states (**a**,**d**), we plot the drop in *CO*_*2*_ concentration within each compartment, the concentrations in the upper and lower cuvettes, and the direction and relative magnitudes of the fluxes into or out of the leaf (**b**,**c**), in comparison to the upper and lower internal concentrations and drop calculated based on a single compartment model (**c**,**f**). In the initial state (**a**), the actual Δ*c*_*i*_’s for the compartments of 4 and 10 PPM respectively (**b**) are far below the single compartment model Δ*c*_*i*_ estimate of 35 PPM (**c**), consistent with the over-estimate of *R*_*ias*_ by that model. In the final state (**d**), the high resistance compartment is nearly isolated from the lower cuvette, which tracks the high conductance compartment as it maintains zero net flux (**e**). The surface resistance of the upper surface as a whole is high enough that the single compartment model underestimates *c*_*i*_ at the upper surface for the high conductance compartment, resulting in an apparent gradient inversion (**f**).

## Notes

### Competing Interest Statement

The authors have declared no competing interest.

