## Supplemental Information Notes S1 for "Ohmic analogies and metaphorical circuits: Vascular partitioning of leaf air spaces and stomatal patchiness can create apparent undersaturation and gradient inversion"

### $CO_2$ gradients in an amphistomatous leaf with asymmetric fluxes

*F. E. Rockwell*[1]

March 14, 2024

#### 1 Diffusion with homogenous assimilation

Consider a one dimensional domain in the z direction, with fluxes of $CO_2$ at both the top (z=0) and bottom (z=L). With an assumption that assimilation is homogenous though the leaf thickness, in steady state, conservation of $CO_2$ leads to a governing equation for the mol fraction of $CO_2$ $\chi$ as,

$$\frac{\partial \chi(z)}{\partial t} = \frac{\phi c D}{1.6\tau}\frac{\partial^2 \chi(z)}{\partial z^2} - A_v = 0 \qquad A_v = \frac{A}{L}. \tag{1}$$

Non-dimensionalizing (1) motivates a change in variables to,

$$\frac{\phi c D}{1.6\tau}\frac{\Delta\chi}{L^2}\frac{\partial^2 \xi(Z)}{\partial Z^2} = A_v, \qquad \xi(Z) = \frac{\chi(z)}{\Delta\chi},\ Z = \frac{z}{L}. \tag{2}$$

The non-dimensional governing equation specifies the choice of $\Delta\chi$,

$$\frac{\partial^2 \xi(Z)}{\partial Z^2} = 1.6\frac{A_v \tau L^2}{\phi c D \Delta\chi}, \qquad \Delta\chi = 1.6\frac{A_v \tau L^2}{\phi c D}. \tag{3}$$

Integration of (3) leads to the form of the solution,

$$\frac{\partial^2 \xi(Z)}{\partial Z^2} = 1 \quad \rightarrow \quad \frac{\partial \xi(Z)}{\partial Z} = Z + a_1 \quad \rightarrow \quad \xi(Z) = \frac{Z^2}{2} + a_1 Z + a_2. \tag{4}$$

The boundary conditions equate the fluxes in the leaf interior at each epidermal surface to the transpiration flux at the external surface, calculated as the mole fraction difference across the surface times the conductance of that surface,

$$-\frac{\phi c D}{\tau}\frac{\partial \chi(z)}{\partial z}\bigg|_{z=0} = g_u(\chi_u - \chi(0)), \tag{5}$$

$$-\frac{\phi c D}{\tau}\frac{\partial \chi(z)}{\partial z}\bigg|_{z=L} = g_l(\chi_l - \chi(L)), \tag{6}$$

where $\chi_u$ and $\chi_l$ are the mole fractions in the upper and lower cuvette's respectively. Here the sign convention is that fluxes that are downward in the positive z direction are positive. Non-dimensionalizing the boundary conditions leads to,

$$-\left.\frac{\partial\xi(Z)}{\partial Z}\right|_{Z=0} = \beta_u(\xi_u - \xi(0)), \quad \beta_u = \frac{g_u \tau L}{\phi c D}, \tag{7}$$

$$-\left.\frac{\partial\xi(Z)}{\partial Z}\right|_{Z=1} = \beta_l(\xi(1) - \xi_l), \quad \beta_l = \frac{g_l \tau L}{\phi c D}, \tag{8}$$

Here we have found two Biot numbers $\beta_u$, $\beta_l$ that represent the balance of each surface conductance to the conductance across the intercellular airspace (IAS).

Applying the upper surface boundary condition (7) to the solution (4) finds that,

$$-\left.\frac{\partial\xi(Z)}{\partial Z}\right|_{Z=0} = -a_1, \text{ and } \xi(0) = a_2 \quad \rightarrow \quad a_1 = \beta_u a_2 - \beta_u \xi_u. \tag{9}$$

Applying the lower surface boundary condition (8) to the solution (4) leads to,

$$-\left.\frac{\partial\xi(Z)}{\partial Z}\right|_{Z=1} = -1 - a_1, \text{ and } \xi(1) = \frac{1}{2} + a_1 + a_2 \tag{10}$$

$$-1 - a_1 = \beta_l\left(\frac{1}{2} + a_1 + a_2\right) - \beta_l \xi_l \tag{11}$$

Combining (9) and (11) leads to a solution for the second integration constant,

$$a_2 = \frac{\beta_u \xi_u + \beta_l \xi_l - \beta_l/2 + \beta_l \beta_u \xi_u - 1}{\beta_u + \beta_l + \beta_l \beta_u} \tag{12}$$

Replacing dimensions, the complete solution has the form,

$$\chi(z) = \Delta\chi\left(\frac{z^2}{2L^2} + a_1 \frac{z}{L} + a_2\right), \tag{13}$$

$$\Delta\chi = 1.6\frac{A\tau L}{\phi c D}, \tag{14}$$

$$a_1 = \beta_u\left(a_2 - \frac{\chi_u}{\Delta\chi}\right), \tag{15}$$

$$a_2 = \frac{\beta_u \frac{\chi_u}{\Delta\chi} + \beta_l \frac{\chi_l}{\Delta\chi} - \beta_l/2 + \beta_l \beta_u \frac{\chi_u}{\Delta\chi} - 1}{\beta_u + \beta_l + \beta_l \beta_u}, \tag{16}$$

$$\beta_u = \frac{g_u \tau L}{\phi c D}, \tag{17}$$

$$\beta_l = \frac{g_l \tau L}{\phi c D}. \tag{18}$$
